# Sequencing Micronuclei Reveals the Landscape of Chromosomal Instability

**DOI:** 10.1101/2021.10.28.466311

**Authors:** Catalina Pereira, Ana Rita Rebelo, Dashiell J. Massey, John C. Schimenti, Robert S. Weiss, Amnon Koren

**Affiliations:** Department of Biomedical Sciences, Cornell University, Ithaca, New York 14853, USA; Department of Molecular Biology and Genetics, Cornell University, Ithaca, New York 14853, USA

## Abstract

Genome instability (GIN) is a main contributing factor to congenital and somatic diseases, but its sporadic occurrence in individual cell cycles makes it difficult to study mechanistically. One profound manifestation of GIN is the formation of micronuclei (MN), the engulfment of chromosomes or chromosome fragments in their own nuclear structures separate from the main nucleus. Here, we developed MN-seq, an approach for sequencing the DNA contained within micronuclei. We applied MN-seq to mice with mutations in *Mcm4* and *Rad9a*, which disrupt DNA replication, repair, and damage responses. Data analysis and simulations show that centromere presence, fragment length, and a heterogenous landscape of chromosomal fragility all contribute to the patterns of DNA present within MN. In particular, we show that long genes, but also gene-poor regions, are associated with chromosome breaks that lead to the enrichment of particular genomic sequences in MN, in a genetic background-specific manner. Finally, we introduce single-cell micronucleus sequencing (scMN-seq), an approach to sequence the DNA present in MN of individual cells. Together, sequencing micronuclei provides a systematic approach for studying GIN and reveals novel molecular associations with chromosome breakage and segregation.

## Introduction

Genome instability, a hallmark of cancer, results from various intrinsic and extrinsic sources of genotoxic stress. GIN does not manifest uniformly across the genome; instead, certain genomic regions and features are more prone to one or more forms of chromosome damage, breakage, and rearrangements. An important manifestation of this genomic heterogeneity are fragile sites: specific regions that are prone to breakage and rearrangements and that are cytologically observed as gaps or constrictions on metaphase chromosomes. Fragile sites can be categorized as common or rare, depending on their frequency in the population, and are typically observed following induction using mild applications of replication inhibitors (e.g. Aphidicolin). Late-replicating genomic regions are particularly prone to being fragile. However, while late replication is generally associated with a low gene density, numerous studies have implicated very long genes as potent drivers of fragile sites (1–8). Another category are early-replicating fragile sites (ERFS), which tend to form due to replication-transcription conflicts at gene-rich, early-replicating genomic regions (9).

Identification of fragile sites or otherwise common sites of GIN is informative regarding their mechanisms of formation. However, mapping chromosomal fragility in an accurate, comprehensive genome-wide scale poses technical challenges. One approach is to analyze chromosomal rearrangement in tumor genomes. This accurately reveals past chromosomal breakage events, but only those that have been selected during cancer evolution rather than the full distribution of fragility events. Alternatively, several genomic assays have been developed to directly map the sites of chromosomal double strand breaks (10, 11), however these assays are prone to background noise that obscures the signal originating from less common events.

Another approach for mapping the landscape of chromosome fragility is to utilize one of the direct cellular outcomes of such instability. Specifically, broken, mis-segregated or unreplicated chromosomes or chromosome fragments can become separated from the main nucleus at mitosis and result in the formation of micronuclei (MN). MN are a common manifestation of genomic instability, are known to undergo partial and abnormal replication, and have the potential to be re-incorporated into the primary nucleus after undergoing extensive rearrangement. Micronuclear DNA can give rise to chromothripsis, which is observed in numerous solid cancer types (12, 13), or to the spillage of DNA into the cytosol, where it can elicit the cGAS/STING inflammatory response (14); both events contribute to tumorigenesis.

We and others have shown that the level of MN is a reliable readout of genomic instability (15–20). Specifically, anucleated red blood cells (RBCs) lose their primary nucleus during terminal differentiation but retain micronuclei that may have formed during prior mitotic cell divisions. In mice, these micronucleated erythrocytes are not efficiently cleared from the circulation and can thus be analyzed from peripheral blood samples. Quantifying the levels of MN-containing red blood cells provides a means of assessing the rate of genomic instability in various mouse mutants and under different genotoxic stresses. Moreover, identifying the genomic regions giving rise to the DNA fragments that get trapped inside micronuclei can reveal preferential sites of chromosomal instability, which themselves may vary in different genetic strains and conditions. Indeed, a previous study sequenced the DNA in RBC MNs, revealing the preferential mis-segregation of acentromeric chromosomal fragments as well as sites of elevated DNA breakage (21). Here, we further developed a similar approach of sequencing MN DNA from cell populations as well as from single cells, and applied it to wild type (WT) mice and two mutant mouse models with different types of genomic instability. Data analysis and simulations link MN formation to centromere presence and to a heterogenous landscape of chromosomal fragility related to long genes and to gene-poor regions.

## Results and Discussion

To study the DNA sequences harbored within MN, we used flow cytometry to isolate RBCs that contained a measurable amount of DNA above background, indicative of MN presence. We specifically isolated normochromatic erythrocytes (NCEs), which harbor spontaneous micronucleation events that happen during erythropoiesis. These cells have a lifespan of approximately 120 days and provide a readout of steady state levels of MN. The precursors of NCEs are reticulocytes, which are short-lived products of the most recent mitotic cycle and as such can be used to quantify MN after extrinsic or transient sources of genotoxic stress (Figure 1; (15, 16, 19, 20)).

**Figure 1.**
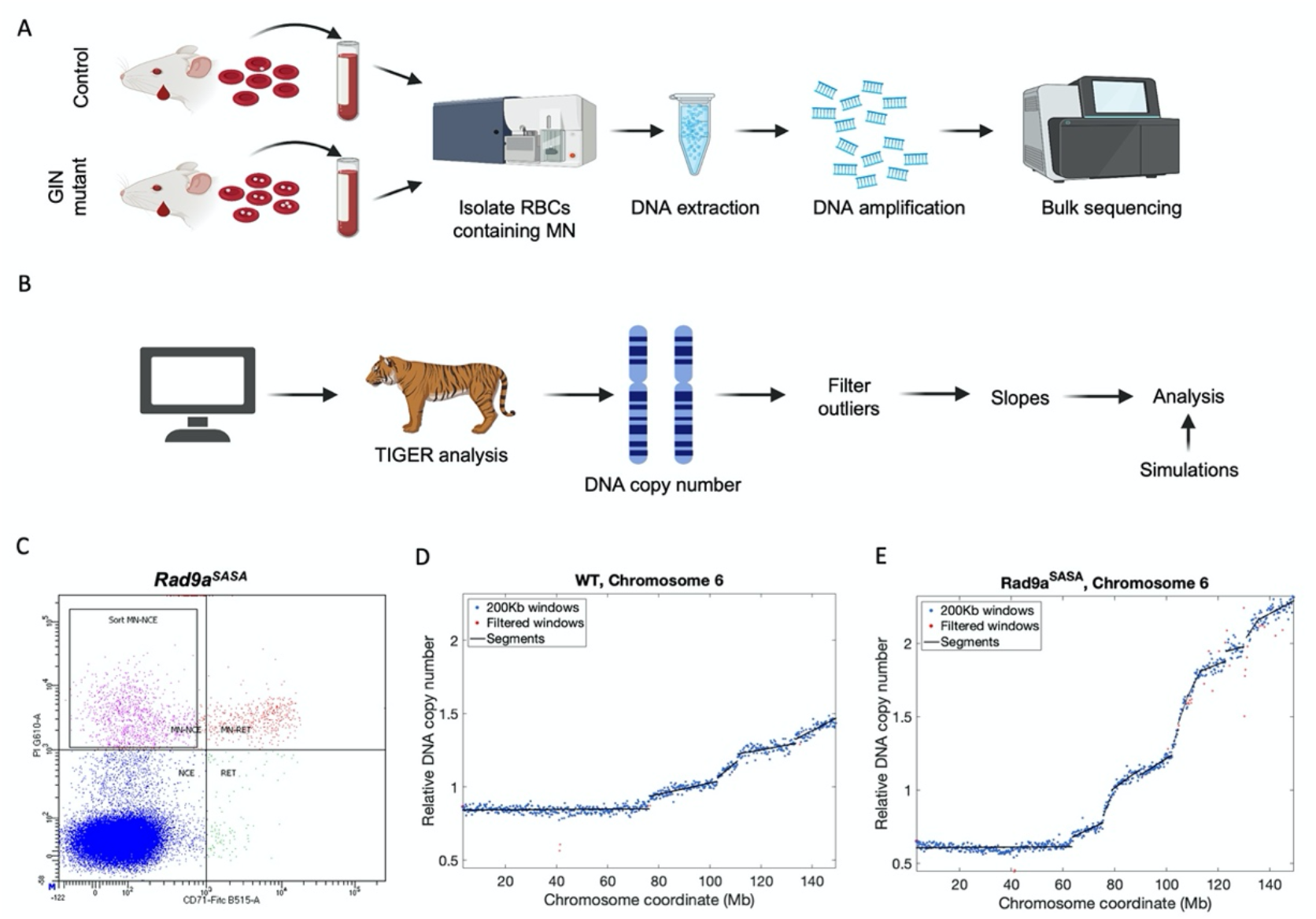
MN-seq. **A.** Experimental pipeline. Peripheral blood is flow-sorted to isolate MN-containing RBCs, DNA is extracted, amplified and sequenced. **B.** Analysis pipeline. TIGER (23) is used to infer DNA copy number across chromosomes, outliers (that result from technical noise or from copy number variation (CNVs)) are removed, and chromosomes are segmented into regions with continuous copy number states from which slopes are calculated. In parallel, computer simulations are used to predict the patterns of MN DNA resulting from different scenarios of chromosome breakage and chromosome fragment segregation into micronuclei. The slope data and simulations are jointly considered for interpreting the data and analyzing the properties of genomic instability. (Cartoons in panels A and B were created with BioRender.com) **C.** Isolation of MN-containing cells by flow sorting of DNA-containing (PI-positive) normochromatic erythrocytes (NCEs), which have lost expression of the transferrin receptor CD71. Reticulocytes (RET, upper right quadrant) retain CD71 expression. **D-E.** Examples of MN-seq data of chromosome 1 in WT (**D**) and *Rad9a^SA^* (**E**) strains. Each blue dot represents the relative DNA copy number within a 100Kb window of uniquely alignable sequence. Copy number is normalized to a genome-wide mean of 1. Red dots have been filtered-out as outliers or suspected CNVs. Black lines represent continuous segments, from which the slopes are calculated. In these examples, DNA copy number initially decreases and then sharply increases in several discrete shifts that are more pronounced upon mutation of *Rad9a*.

MN were isolated from two GIN mouse models and their respective WT strains. The first was the *Mcm4^chaos3^* (*Chaos3*) strain, originally identified via a screen for mice with elevated MN (15), that harbors a mutation in *Mcm4* (22), part of the Mcm2-7 complex component of the replicative helicase. The second model were mice harboring a mutation in *Rad9a*, part of the RAD9A-RAD1-HUS1 (9A-1-1) clamp complex that recruits DNA damage signaling and repair factors to damage sites and functions as part of the DNA damage checkpoint response. Specifically, we mutated a phosphorylation site in the RAD9A C-terminal tail (S385A) that is essential for interaction with TOPBP1 and 9-1-1 mediated ATR activation (hereafter referred to as the *Rad9a^SA^* mutant). As expected, *Chaos3* mice had a 20.9-fold increase in cells containing MN-NCE compared to isogenic WT mice. Homozygous *Rad9a^SA^* mice had an average of 8.7-fold increased rate of micronuclei relative to control mice (Table S1). We sorted between 83,000 and 800,000 cells per strain, isolated and amplified DNA and performed whole-genome sequencing (see Methods). We refer to this approach as MN-seq.

To analyze the genomic regions present in MN, we modified an analytic pipeline to call DNA copy number along chromosomes (23) (Figure 1B; Methods). Briefly, sequencing reads were counted in windows of 200Kb of uniquely alignable sequence and corrected for GC content effects on read coverage. Copy number segmentation and filtering was used to remove outlier data points as well as short segments that were grossly inconsistent with their surrounding regions. Subsequently, the slope of the filtered copy number values in each segment were calculated (Figure 1D-E; Table S2).

Analysis of the resulting changes in DNA representation along chromosomes revealed a complex landscape of genomic regions giving rise to MN DNA. At a broad scale, micronuclei tended to harbor increasing amounts of DNA from the telomeric ends of chromosomes (Figure 1D-E, Figure 2, Figure S1). This is likely due to acentric chromosome fragments resolving into micronuclei. This, and the apparently opposite trend at the beginning of chromosomes 1 and 2, both confirm previous observations (21). However, we also noticed many significant changes in the steepness of copy number slopes across the chromosomes, including both increases and decreases. There were also additional examples (besides chromosomes 1 and 2) of negative slopes, in which MN DNA is less represented from the proximal to distal parts of a chromosomal region. More generally, the level of MN varied by up to ~12-fold across the genome. Notably, all these patterns showed variations between WT mice, *Rad9a^SA^* mutants, and *Mcm4^Chaos3^* mice, while they were highly reproducible in repeat experiments of the same genetic background (Figure 2; Figures S1-S2). Differences in MN levels between mouse strains and sexes have been reported before (20, 24). Although our sample size is not large enough to ascertain sexspecific differences in MN DNA patterns, we did notice a highly elevated rate of MN fragments towards the right end of the X chromosome in females, in both WT strains and even more so in GIN mutants; this could conceivably be explained by a high level of chromosomal instability of the inactive X chromosome in female samples.

**Figure 2.**
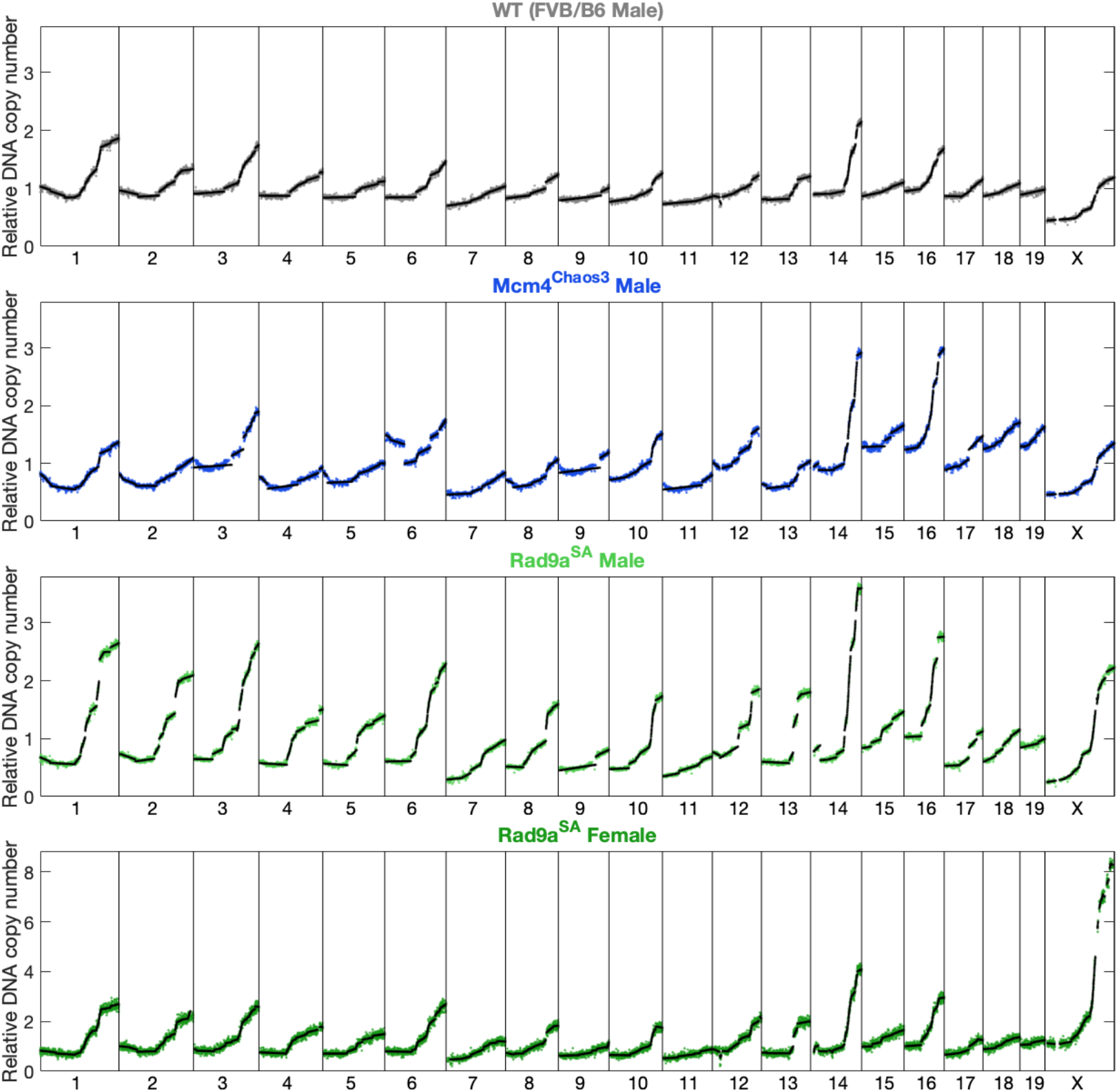
The landscape of micronuclei DNA across the genome of different mouse strains. As in Figure 1D-E, for all chromosomes and various strains analyzed in this study (see Figure S1 for the remaining samples). Several notable patterns are observed, including negative and positive slopes, slopes of different steepness, slope shifts (both upward and downward), and differences between strains. The patterns are more subtle in WT than in mutant mice, and are different between *Mcm4^Chaos3^* and *Rad9a^SA^* mutants strains. Also note the different DNA copy number scale for the *Rad9a^SA^* female sample, which reflects the extreme MN levels towards the right end of the X chromosome. The WT sample is a male FVB/B6 (the control strain for the *Rad9a^SA^* mutants).

We confirmed that sharp shifts in DNA copy number, for example the one on chromosome 6 of *Mcm4^Chaos3^* mice, do not result from copy number variation in the genome itself of these strains (Figure S3). We cannot, however, rule out the possibility that structural rearrangements such as inversions or translocations influence the observed MN copy number patterns.

To interpret the various patterns of MN DNA representation, we used *in silico* simulations. Two main parameters hypothesized to affect MN DNA were simulated: the locations of breaks along a chromosome, and which part of the chromosome results in a micronucleus after breakage. We considered random locations of breaks uniformly distributed along the lengths of chromosomes, regional changes (increases or decreases) in the rate of breaks in different parts of a chromosome, and hotspots comprising sharp elevations of chromosome break rates at specific locations. With regards to the fragments that are retained within MN, we considered whether the acentric or, reciprocally, centric fragments of a chromosome preferentially resolve into micronuclei. We also considered the possibility that the length of a fragment, rather than the presence of a centromere, is the determining factor for inclusion in MN. We thus simulated chromosome breaks in individual chromosomes, followed by aggregating many such hypothetical chromosomes in order to obtain MN DNA content patterns that could be compared to the empirical MN-seq data.

Simulations confirmed that if acentric chromosomal fragments are those that tend to result in MN, a gradual increase in DNA representation will be observed along the length of the chromosome (maintaining the assumption that the centromeres are at the left end). This increase will be gradual if break sites are random (Figure 3B), but will accelerate in regions of high breakpoint density (Figure 3G), decelerate in regions of low breakpoint density (Figure 3H), and undergo an abrupt shift if there are localized hotspots of chromosomal breakage (Figure 3E). On the other hand, if centric fragments are those that compose MN DNA, we would expect to see a gradual *decrease* in DNA content along a chromosome (Figure 3C). If centric and acentric fragments are equally likely to segregate into MN, a flat pattern of DNA content is expected to emerge (Figure 3D), which would be difficult to discriminate from large regions with low break rates (Figure 3H). Last, if the main determinant of fragment capture in MN is the length of the fragment (in addition to, or instead of, centromere presence), then the pattern of MN DNA could be expected to show a decrease followed by an increase along the chromosome (or *vice versa* in the less likely scenario in which long rather than short fragments tend to be captured in MN). In such cases, the length preference (and potentially association with centromere presence) will determine the site of slope shift. A specific case would be such a length dependence together with a spike in break rate at approximately the same inflection point – this scenario can explain the unusual pattern seen in *Mcm4^Chaos3^* MN on chromosome 6 (Figure 2; Figure 3F). We also considered the possibility of two breaks occurring on the same chromosome, with the interstitial chromosome fragment either preferentially retained in MN, or retained at random. In either case, the resulting MN patterns did not resemble the observed data (Figure S4). This is also consistent with the relatively modest rate of MN across the studied strains (<5% of cells per sample; Table S1), which would predict that two breaks occurring on the same chromosome in the same cell would be rare.

**Figure 3.**
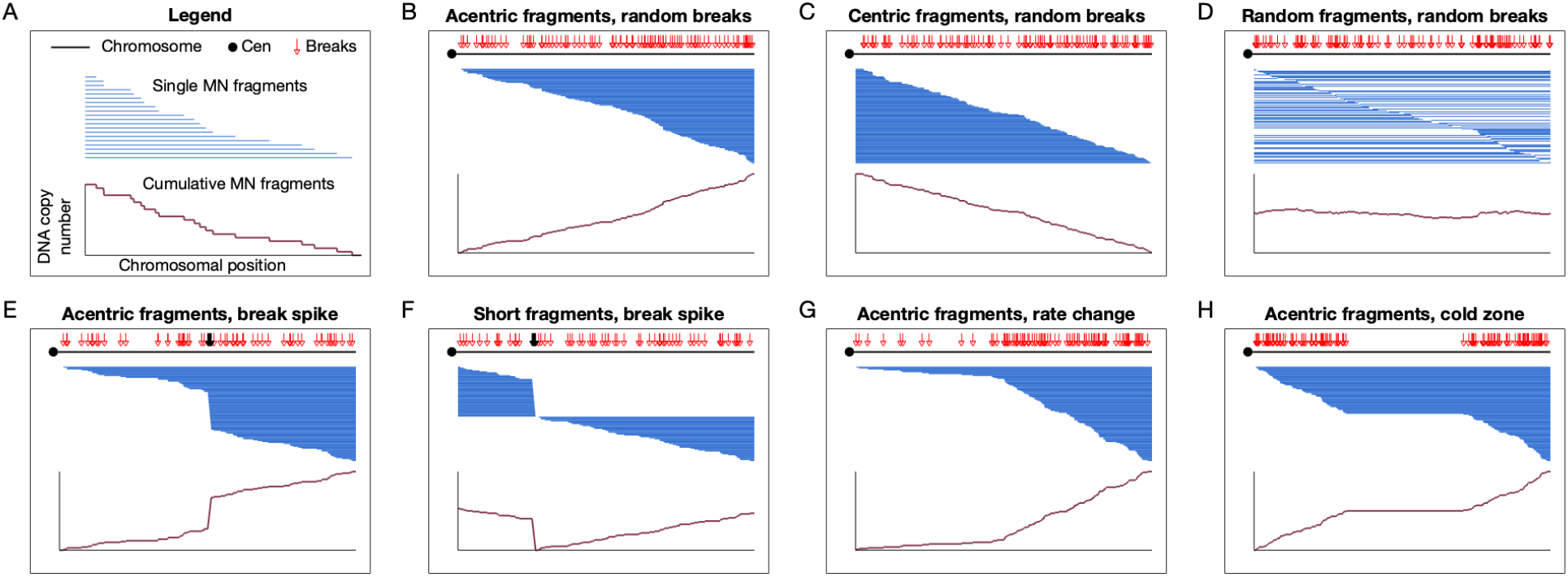
Simulations of MN DNA representation as a function of chromosomal break sites and preference for retained fragments. **A.** Explanation of the visual representation of simulated MN DNA fragments. We simulated an artificial chromosome of 10Kb (black line) with the centromere on the left side (circle). We then introduced 100 chromosome breaks (red arrows) at either random locations or at variable rates along the chromosome. We simulated scenarios in which, after a chromosome is broken, the centric, acentric, or a randomly chosen fragment is retained in a micronucleus. Blue lines represent individual fragments that would be retained following a given break (20 are shown in A, and one under each corresponding chromosomal break in other panels). The cumulative MN fragment graph (bottom) represents the count of single DNA molecules covering regions along the chromosome, and corresponds to the expected data observed from MN-seq analysis. **B.** Random breaks with only acentric fragments retained in micronuclei. **C.** Random breaks with only centric fragments retained. **D.** Random breaks with either centric or acentric fragments, assigned randomly, retained in micronuclei. **E.** Simulation of a spike with elevated chromosome break rates at the middle of the chromosome; acentric fragments retained. **F.** A breakage spike at 1/4 of the chromosome length, with centric fragments up to that point (i.e. shorter than a 1/4 of the chromosome length) retained, and acentric fragments retained otherwise. **G.** Break rate doubles after the center of the chromosome; acentric fragments retained. **H.** A region in the center of the chromosome devoid of breaks; acentric fragments retained.

Overall, simulations provide plausible explanations (albeit not direct evidence) for the various patterns of MN DNA observed across chromosomes and strains (Figure 2). Having established at least one possible biological interpretation for each of the various patterns of MN DNA content observed in the data, we could now turn to a systematic quantification of the chromosome breakage landscape.

First, we combined all slopes across the genome in each strain separately as well as across all strains together, and analyzed the distributions of their steepness. While most slopes represented a baseline break rate, there was also a mode of segments with negative slopes, a larger mode comprised of slopes with increased steepness that represent regions with elevated chromosomal breaks, and yet another disperse mode with very steep slopes (Figure 4, top). While most negative slopes were attributed to *Chaos3* mice, *Rad9a^SA^* mutants contributed the majority of slope fragments with elevated steepness (Figure 4, bottom). This confirms the strain-specific differences in the chromosomal distribution of break sites and the tendency of different chromosomal fragments to end up within micronuclei.

**Figure 4.**
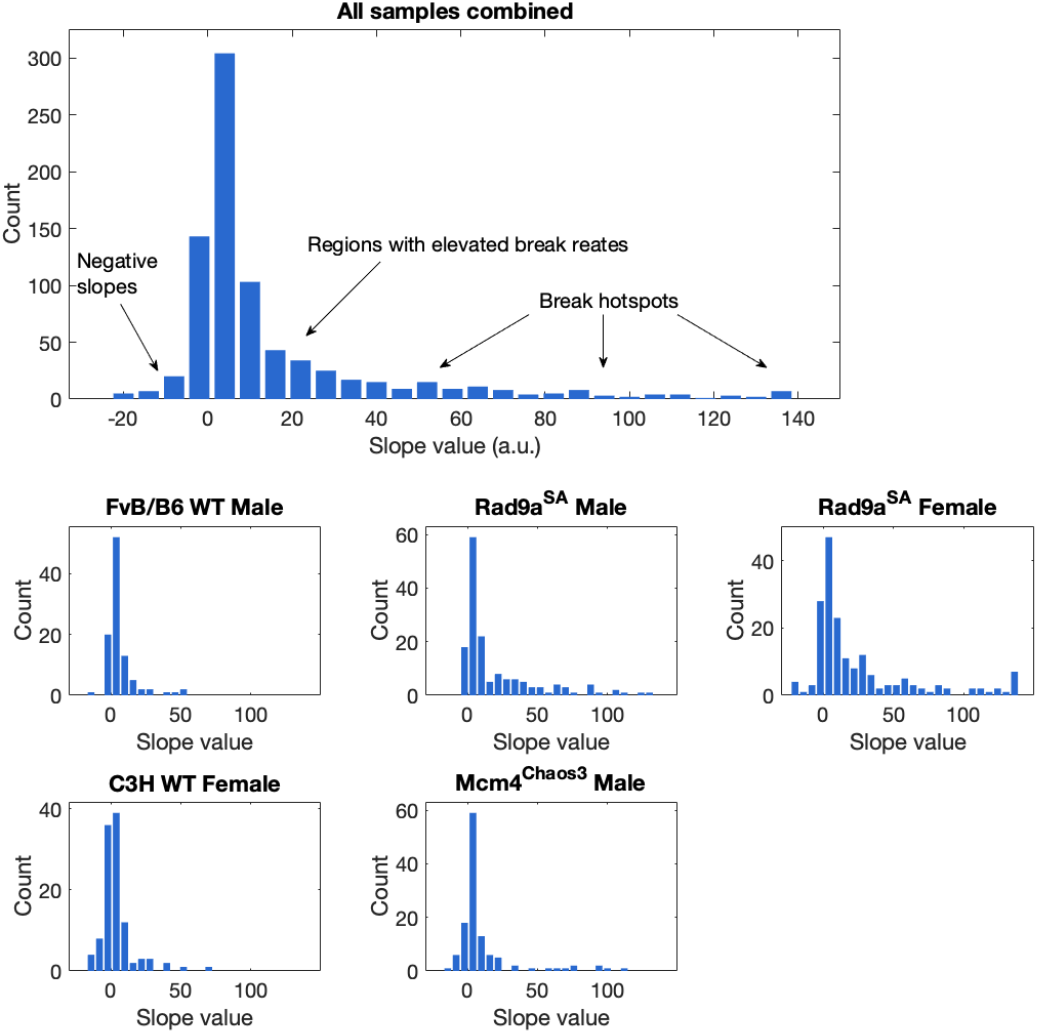
Statistics of MN-seq slopes. Slopes were combined across the genome and, in the top panel, across all samples (sample *Mcm4^Chaos3^* female had lower sequence coverage and was not included in these analyses and in the figure; see Methods). The distribution of slope values is shown. Bottom panels show individual samples; only one repetition for *Rad9a^SA^* male is shown.

We next analyzed the slopes in individual chromosomes and strains and compared them to the locations of genes, DNA replication timing, and early-replicating fragile sites mapped in mouse. Visually, steep slopes appeared to map to late-replicating, gene-poor regions, while sharp shifts between slopes often mapped near long genes (Figure 5). For example, the sharp drop on chromosome 6 in Chaos3 mapped within ~200Kb from *CNTNAP2*, the second-longest gene in the mouse genome (2.24Mb) and a known fragile site in humans and mice (1, 25). Other breakpoints with sharp shifts in DNA copy number between segments mapped to the known fragile sites associated with *LRP1B* (2.06Mb), *MACROD2* (1.99Mb), and *FHIT* (fragile histidine triad gene; 1.57Mb), which contains FRA3B, the most active common fragile site in many human cell types (1, 4, 26, 27). Overall, 9 of the 10 longest genes in the mouse genome were within 0.5Mb from a slope shift breakpoint in DNA copy number segments.

**Figure 5.**
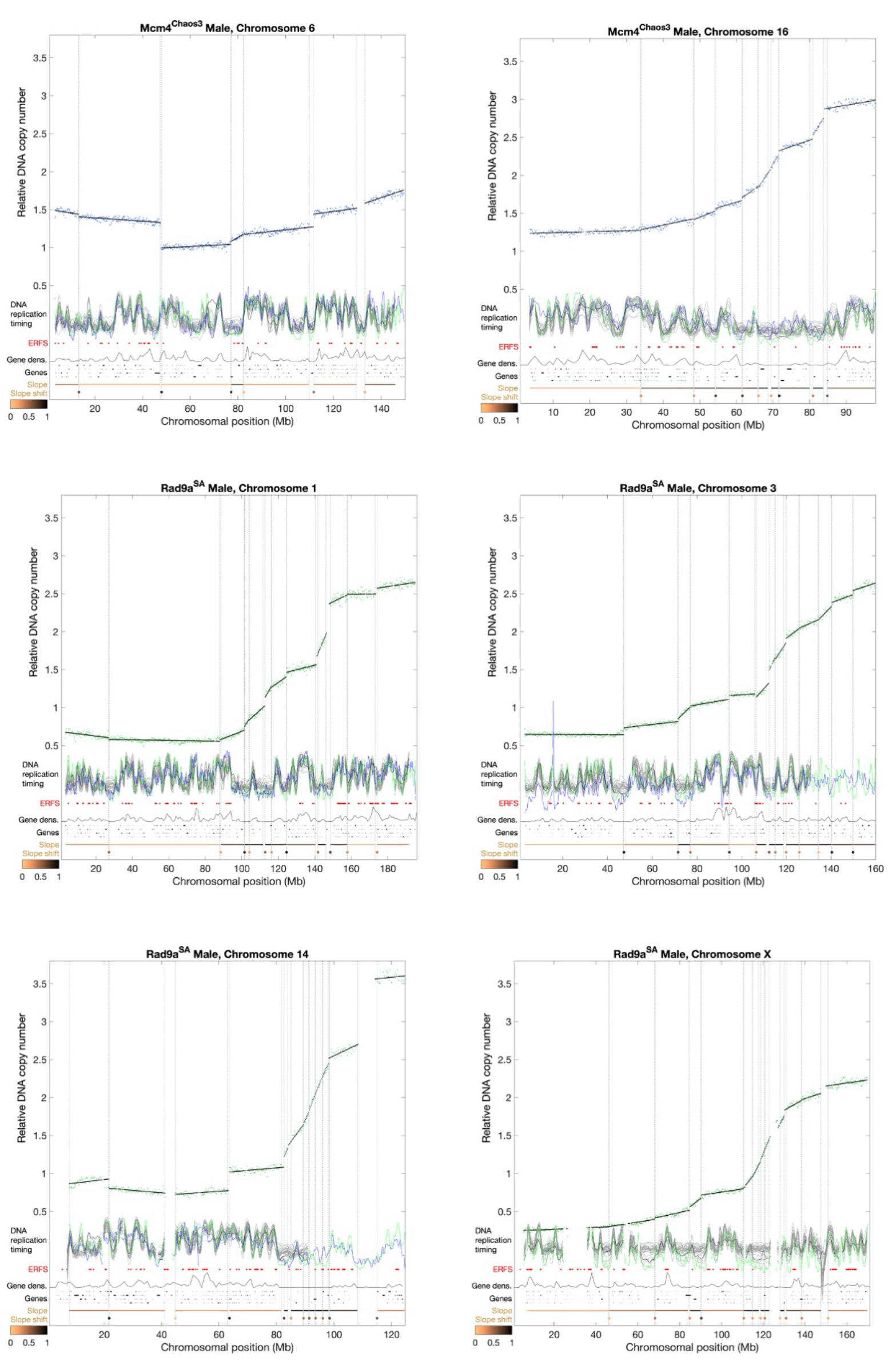
Examples of MN-seq data and slopes in individual chromosomes and samples together with other relevant genomic data. Dashed grey lines mark the borders of slope segments. Note that steep slopes are over-sensitive to being fragmented into several segments that may have similar slopes, thus breakpoints along steep slopes may represent the slopes themselves rather than discrete shifts as seen elsewhere. DNA replication timing: replication profiles measured from 29 different cell types (grey; (28)), pre-B cells (black; (28)), MEFs (green; (23)) and mouse pluripotent stem cells (blue; (23)). Higher values represent earlier replication. Note that some replication timing data is missing for chromosome 14 (due to LiftOver) and chromosome X (due to sex of samples). ERFS: early replicating fragile sites (9). Gene density: the number of genes in 1Mb windows. Genes: individual genes (shown on four vertical levels). Slopes and slope shifts are represented with a color code (colormap underneath) with orange representing shallow slopes and subtle shifts, and darker shades representing steeper slopes and more extreme break rate shifts.

Statistical analysis of these associations confirmed that, genome wide and across strains, there was a strong relationship between steep slopes and both low gene density and late replication (Figure 6). These associations followed an exponential fit rather than a linear relationship, with the majority of steep slopes occurring in regions with less than one gene per megabase (compared to a genome-wide mean of seven genes/Mb; Figure 6A) and with replication timing at least one standard deviation later than the mean (Figure 6B). These associations were observed also in individual chromosomes (Figure S5). Segment breakpoints were also associated with late replication, although not as strongly as slope steepness (Figure 6C). On the other hand, there was a compelling association between breakpoints and long genes, with many segment borders falling close to some of the longest genes in the genome (Figure 6D). Overall, of the 3304 genes within 0.5Mb from a breakpoint, 55 were longer than 0.5Mb, compared to 148 genes in the genome of this length out of a total of 21,958 genes (chi square p = 2.15×10^−9^). We did not find a significant association between breakpoints and ERFS (Figure 5).

**Figure 6.**
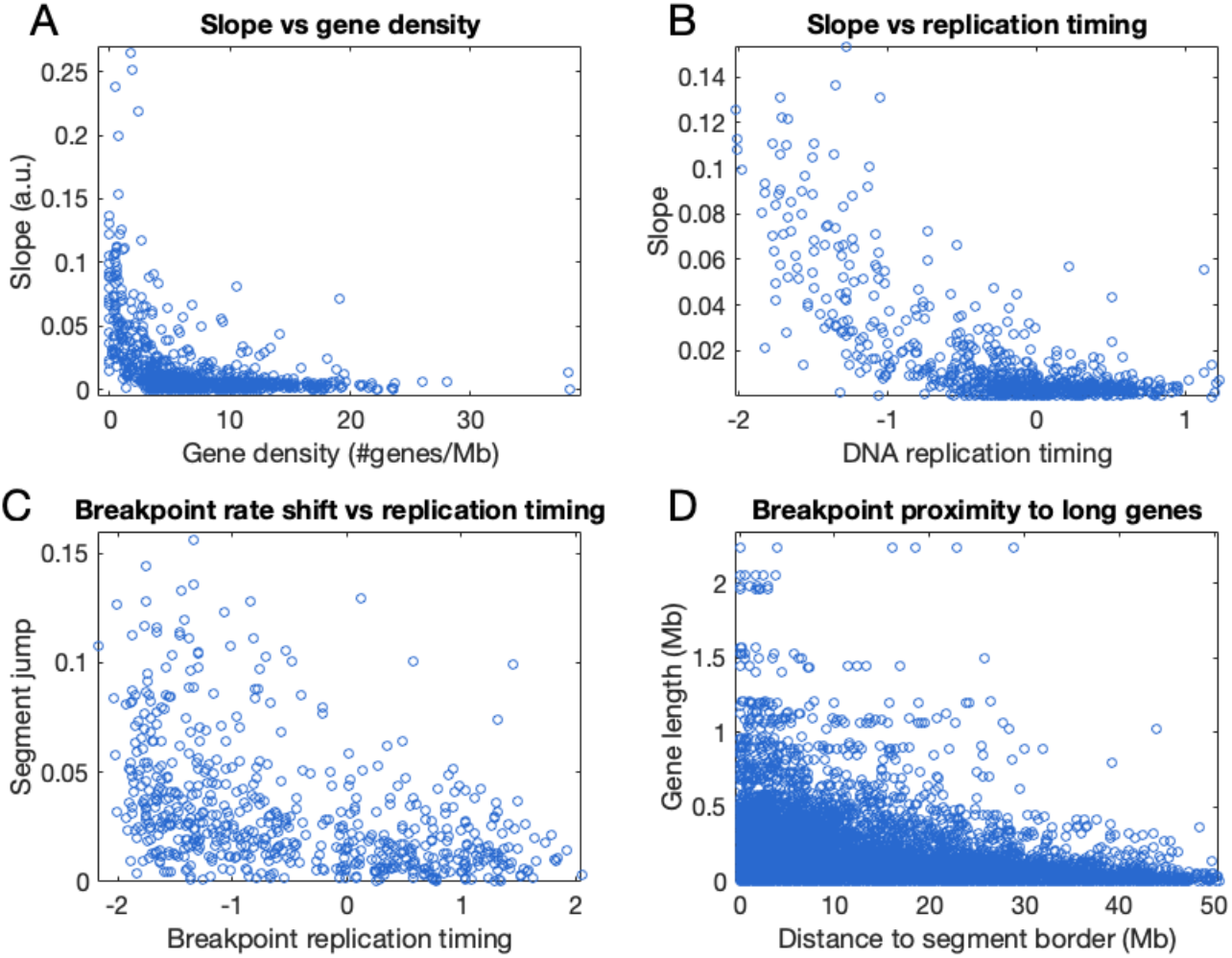
Global correlations of break rates with other genomic features. **A.** Segment slope as a function of gene density. **B.** Segment slope as a function of DNA replication timing in MEFs. Replication timing is presented in units of standard deviation from the mean, which is set to zero across the genome. Negative values represent late replication and positive values represent early replication. Replication profiles from other cell types provided similar results. **C.** Segment shift magnitude as a function of DNA replication timing in MEFs. **D.** Gene length as a function of the distance of a gene to the nearest segment breakpoint. Calculated for all breakpoint-gene pairs. Sample *Mcm4^Chaos3^* female was not used in any of the analyses.

While an association between long genes and chromosome fragility is well-established (1–8), our results point to an independent contribution of gene-poor chromosomal regions to chromosome breakage. Gene-poor regions were also recently shown to associate with chromosomal breaks in human embryos (29). Notably, the locations of both long genes and gene-poor regions are correlated with late replication. However, they manifest on genome stability in different ways: long genes tend to be associated with strong breakage hotspots, while gene density has a more subtle albeit more comprehensive association with the overall background rates of breaks along chromosomes. While long genes may cause chromosome breaks through transcription intermediates, it is less clear how gene-poor regions could relate to broad chromosome stability. Gene density is also a highly heterogenous feature of chromosomes and cannot by itself predict the rates of MN formation observed in our data, thus additional factors likely interact to shape the genomic landscape of fragility. Further improvement in data resolution will enable finer mapping of chromosomal break rates as well as the identification of break hotspots at higher resolution.

Another limitation of MN-seq is that it relies on sequencing DNA from many MN-containing cells. The analyzed signal therefore derives from the superimposition of many events, making it difficult to infer the molecular events that gave rise to individual micronuclei (for example see Figure 3). However, chromosomal breaks are expected to happen at different locations in different cells, thus ensemble approaches (such as MN-seq and other genomic approaches) represent the cumulative fragility landscape yet miss individual events that are diluted by background signals from other cells. Recently, several approaches have been developed for whole-genome sequencing of DNA from single mammalian cells (30–34). These approaches rely on isolating individual cells, extracting and barcoding the DNA from each cell, and then pooling, amplifying, and sequencing the DNA. We reasoned that a similar approach could be applied to MN-containing erythrocytes, in particular since the cells are still intact and can be isolated. While the amount of DNA in a single micronucleus is very small, the signal that is sought is binary, i.e. the presence or absence of DNA representing large chromosomal fragments, making it amenable for such analyses. We hence developed single-cell micronucleus sequencing (scMN-seq), in which the DNA in (one or more) micronuclei contained within individual cells is separately sequenced. We used the 10x Genomics Single-Cell CNV platform for cell isolation, barcoding and DNA amplification, followed by whole genome Illumina sequencing. Although most cells were lost during the process or failed to amplify, we did obtain clear signals from 14 single MN-containing cells from WT, *Rad9a^SA^* and Chaos3 mice. Analysis of the DNA present in these cells revealed that micronuclei (one or more per cell) contained DNA from between one to three chromosomes. In some of the cells, entire chromosomes were present in micronuclei, while in other cases, either an acentric (right) or, less often, centric (left) portion of a chromosome was present in a micronucleus (Figure 7). These observations confirm our interpretation of the bulk MN-seq data as well as the *in silico* simulations (which are fundamentally a simulation of single micronuclei), including that centromere-containing chromosomal fragments can indeed result in micronuclei. More generally, we suggest that the scMN-seq approach, applied on a larger scale and in different genetic background and conditions, is a promising direction to elucidate mechanism of genomic instability and identify specific sites of chromosome breakage and the rules governing which fragments preferentially result in MN formation.

**Figure 7.**
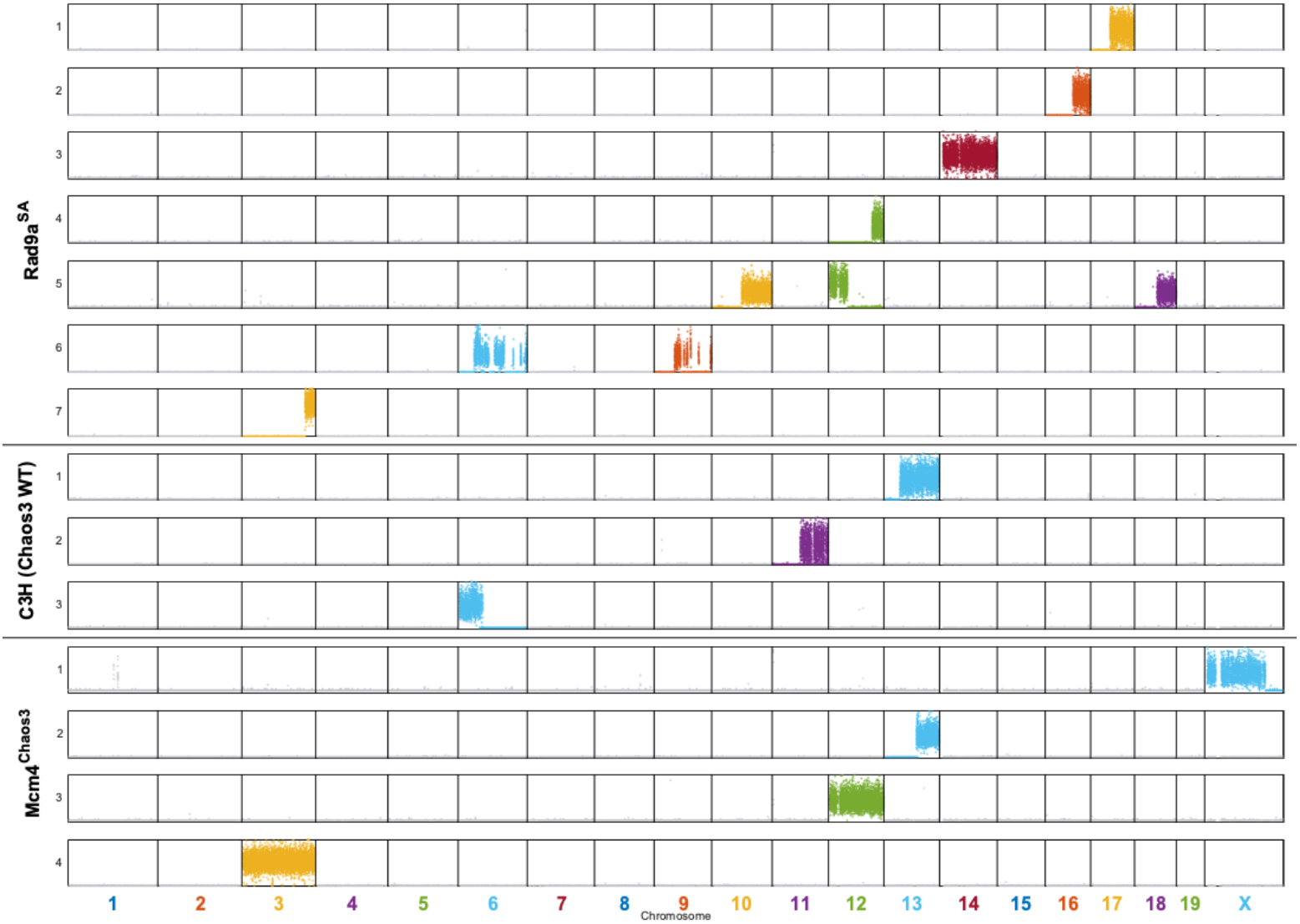
Single micronucleus sequencing. MN-containing NCEs were flow sorted and subjected to microfluidic isolation and library preparation using the 10X Genomics CNV platform, followed by whole genome sequencing. Each row shows the sequence content of MN from a single cell.

To summarize, we’ve shown that micronuclei from mouse peripheral blood red blood cells can be sequenced, in bulk and from individual cells, and that the resulting data is highly informative with regards to the landscape of chromosome breakage. MN-seq and scMN-seq provide informative and straightforward approaches to study genome stability and can be applied in larger scales to many genetic backgrounds, genotoxic stresses and combinations thereof. A main strength of these approaches over alternatives is that they comprise *in vivo* methods for quantifying chromosome fragility genome-wide. A notable limitation is their specificity for anucleated RBCs. It is likely that the DNA sequences contained in micronuclei will differ among cell types, for example due to differences in chromatin structure or cell-type-specific activities of DNA repair pathways or responses to genotoxic stress. While it is feasible to separate micronuclei from main nuclei in other cell types, this is currently difficult to apply in an accurate and large-scale manner. Last, MN-containing erythrocytes are retained in peripheral blood of mice but are cleared more efficiently by the spleen in humans (35). Performing similar experiments in humans will thus require enrichment for reticulocytes, the recruitment of patients post-splenectomy, or working with other, nucleated cell types.

MN-seq revealed a heterogenous landscape of chromosome fragility and chromosomal fragment retention. Long genes and gene-poor regions both emerge as central factors in chromosome fragility, while centromere presence and fragment length influence the fate of broken chromosomes. While common fragile sites (which are essentially re-discovered here in a systematic way) have previously been linked to long, late-replicating genes, MN-seq further provides a quantification of chromosome breakage rates genome-wide, revealing a new association with gene density. The fragility landscape and the tendency of chromosomal fragments to be retained in micronuclei both differ between mouse strains. Since chromosomal breaks can lead to rearrangements and promote cancer, these strain differences are consistent with different genetic backgrounds having different cancer susceptibilities (for example, *Chaos3* mice have a strong tendency to develop sporadic mammary tumors (22)). Further studies are required in order to establish the pathways by which mutations in specific genes involved in the maintenance of genome stability influence the sites of chromosome breakage.

## Methods

### Mice

All mice used for this study were handled following federal and institutional guidelines under a protocol approved by the Institutional Animal Care and Use Committee (IACUC) at Cornell University. The *Chaos3* strain was previously described (15). *Rad9a^SA^* mice contain a serine to alanine mutation at residue 385 generated by CRISPR/Cas9 gene editing as to be described elsewhere and were maintained in a mixed FVB/NJ and C57BL/6J background.

### MN sorting

Blood was collected from the submandibular vein of adult mice aged between 6-12 weeks old. Blood was collected into heparin-containing tubes and was fixed in pre-chilled methanol in −80°C. Blood cells were analyzed for MN presence following protocol described in Balmus *et. al*. (20). Briefly, cells were washed with sodium bicarbonate to remove methanol, stained with anti-CD71 (Fisher #P01275F05) and treated with RNaseA (Sigma #45-R6513). Cells were subsequently washed and stained with propidium iodine (PI; Thermo Fisher #P3566) to visualize DNA. CD71-negative and PI-positive NCEs were isolated using a Sony MA900 cell sorter.

### MN-seq

Following RBC sorting, DNA was extracted using the MasterPure™ Complete DNA and RNA Purification Kit (Lucigen) following the manufacturer’s instructions. DNA was then amplified using QIAseq Ultralow Input Library Kit (Qiagen) following the manufacturer’s instructions. Whole genome sequencing was performed using the Illumina NextSeq500 with 75bp reads. Sequencing reads were converted into non-mapped bam files and marked for Illumina adaptors and duplicate reads with Picard Tools (v1.138) (http://broadinstitute.github.io/picard/) commands ‘FastqToSam’, ‘MarkIlluminaAdapters’, and ‘MarkDuplicates’. Bam files were aligned to mm10 with BWA mem (v0.7.17).

Samples were processed and sequenced in two separate batches. The first batch consisted of the following samples: FVB/B6 male (*Rad9a* WT; 44.14 million reads obtained); *Rad9a^SA^* male repetition 1 (32.23 million reads); *Rad9a^SA^* male repetition 2 (25.17 million reads); and *Chaos3* male (35.4 million reads). The second batch consisted of samples *Rad9a^SA^* female (24.26 million reads); C3H female (*Chaos3* WT; 24.23 million reads); and *Chaos3* female (3.59 million reads).

### Data processing

GC-corrected read depth data for each sample were generated via TIGER (28) using a read length of 100bp for alignability filtering and a bin size of 10Kb. The segment-filtering step utilized for GC correction in TIGER was not implemented (since the copy number across chromosomes is not uniform). Subsequently, every 20 TIGER windows were merged and the TIGER segment_filt function was applied with an R2 value of 0.01 and std threshold of 0.1. Following this filtering step, remaining segments shorter than 5 windows were also removed, as were segments shorter than 20 windows that had either higher or lower copy number values than both of their flanking segments (i.e. represent an inconsistent change of copy number values along chromosomes). For the first and last segments on each chromosome, the latter criterion was replaced by whether the segments were different by more than one 0.3 copies from their adjacent segment. For the second sample batch, since the sequencing coverage was lower compared to the first batch (which resulted in a higher level of noise), a segment filtering std threshold of 0.3 (instead of 0.1) was used. In addition, since the Chaos3 female sample had yet lower coverage than others, we relaxed the R2 value to 0.05, the std threshold to 0.5, and did not apply the filter for segment that were inconsistent with their neighbors. Due to the relaxed data filtering of the Chaos3 female sample, we did not include it in the statistical analysis of slopes (Figures 4 and 6).

### Segment analysis

Data was smoothed as in TIGER, with a smoothing parameter of 10^−19^. For each segment, the smoothed data was used to fit a linear polynomial curve, from which the slope of the segment was derived.

### Simulations

A linear vector the length of 10,000 was used to simulate a chromosome, with 100 random locations initially chosen as break sites. Break sites were then adjusted based on the selected model: for a spike in break rate, 40% of break sites were removed and replaced with breaks clustered in proximity to each other; for a change in break rate, 75% of breaks were removed and replaced with breaks limited to the right half of the chromosome; for a region with no breaks, 150 breaks were simulated at random locations, sorted by position, and break indexes 50-100 were removed. The fragments retained in MN were then simulated as follows: for centromeric fragments, the left end of the broken chromosomes was assumed as retained; for acentric fragments, the right end was retained; for random fragments, either the left or right fragment were randomly retained; for length-dependence of fragment retention, left fragments were retained if they were shorter than a selected length, and otherwise right fragments were retained. The locations of the breaks were plotted on the simulated chromosomes, and the expected MN fragment resulting from each break and retained in MN in the selected model was plotted as a line. To calculate the expected cumulative MN copy number values, the location indexes of all retained fragments were added and plotted.

### External data

Genes and gene density were based on RefSeq genes (mm10). Early-replicating fragile sites were downloaded from Ref. (9). Replication timing data from Refs. (23, 28) was normalized to a zero mean and one standard deviation. Replication timing data for the X chromosome was not available.

### scMN-seq

Sorted samples of each strain were processed using the 10x Genomics Single-Cell CNV kit on the Chromium Controller (10x Genomics, Pleasanton, CA, USA). Data was processed as described (34). Briefly, sequencing reads associated with each cell-specific barcode were counted in 20Kb windows. To account for low-level background, a minimum read threshold was set for each cell at the 10^th^ percentile of per-window read count. A cell was determined to have a micronucleus if at least 10% of any chromosome had read counts above its minimum read threshold. Multiple barcodes associated with identical patterns of micronuclei were observed in *Rad9a^SA^*, and assumed to be a technical artifact.

## Acknowledgements

We thank Adrian McNairn for providing the *Chaos3* mice. This work was supported by the National Institutes of Health (DP2-GM123495 to A.K and R01 HD095296 to RSW), NSF predoctoral fellowship DGE-1144153 (to CP), and seed grants from the Cornell Intercampus Academic Integration Program and from the Cornell Institute of Biotechnology.

## Data Availability Statement

MN-seq and scMN-seq data reported in this study are available from the Sequence Read Archive (SRA) under accession number PRJNA771079.

## References

1. Smith DI, Zhu Y, McAvoy S, Kuhn R. Common fragile sites, extremely large genes, neural development and cancer. Cancer Lett. 2006;232(1):48–57.

2. Sarni D, Sasaki T, Irony Tur-Sinai M, Miron K, Rivera-Mulia JC, Magnuson B, et al. 3D genome organization contributes to genome instability at fragile sites. Nat Commun. 2020;11(1):3613.

3. Brison O, El-Hilali S, Azar D, Koundrioukoff S, Schmidt M, Nahse V, et al. Transcription-mediated organization of the replication initiation program across large genes sets common fragile sites genome-wide. Nat Commun. 2019;10(1):5693.

4. Le Tallec B, Millot GA, Blin ME, Brison O, Dutrillaux B, Debatisse M. Common fragile site profiling in epithelial and erythroid cells reveals that most recurrent cancer deletions lie in fragile sites hosting large genes. Cell Rep. 2013;4(3):420–8.

5. Le Tallec B, Koundrioukoff S, Wilhelm T, Letessier A, Brison O, Debatisse M. Updating the mechanisms of common fragile site instability: how to reconcile the different views? Cell Mol Life Sci. 2014;71(23):4489–94.

6. Ozeri-Galai E, Tur-Sinai M, Bester AC, Kerem B. Interplay between genetic and epigenetic factors governs common fragile site instability in cancer. Cell Mol Life Sci. 2014;71(23):4495–506.

7. Letessier A, Millot GA, Koundrioukoff S, Lachages AM, Vogt N, Hansen RS, et al. Celltype-specific replication initiation programs set fragility of the FRA3B fragile site. Nature. 2011;470(7332):120–3.

8. Ozeri-Galai E, Lebofsky R, Rahat A, Bester AC, Bensimon A, Kerem B. Failure of origin activation in response to fork stalling leads to chromosomal instability at fragile sites. Mol Cell. 2011;43(1):122–31.

9. Barlow JH, Faryabi RB, Callen E, Wong N, Malhowski A, Chen HT, et al. Identification of early replicating fragile sites that contribute to genome instability. Cell. 2013;152(3):620–32.

10. Canela A, Sridharan S, Sciascia N, Tubbs A, Meltzer P, Sleckman BP, et al. DNA Breaks and End Resection Measured Genome-wide by End Sequencing. Mol Cell. 2016;63(5):898–911.

11. Mirzazadeh R, Kallas T, Bienko M, Crosetto N. Genome-Wide Profiling of DNA Double-Strand Breaks by the BLESS and BLISS Methods. Methods Mol Biol. 2018;1672:167–94.

12. Crasta K, Ganem NJ, Dagher R, Lantermann AB, Ivanova EV, Pan Y, et al. DNA breaks and chromosome pulverization from errors in mitosis. Nature. 2012;482(7383):53–8.

13. Zhang CZ, Spektor A, Cornils H, Francis JM, Jackson EK, Liu S, et al. Chromothripsis from DNA damage in micronuclei. Nature. 2015;522(7555):179–84.

14. Galluzzi L, Vanpouille-Box C, Bakhoum SF, Demaria S. SnapShot: CGAS-STING Signaling. Cell. 2018;173(1):276–e1.

15. Shima N, Hartford SA, Duffy T, Wilson LA, Schimenti KJ, Schimenti JC. Phenotype-based identification of mouse chromosome instability mutants. Genetics. 2003;163(3):1031–40.

16. Levitt PS, Zhu M, Cassano A, Yazinski SA, Liu H, Darfler J, et al. Genome maintenance defects in cultured cells and mice following partial inactivation of the essential cell cycle checkpoint gene Hus1. Mol Cell Biol. 2007;27(6):2189–201.

17. Holloway JK, Mohan S, Balmus G, Sun X, Modzelewski A, Borst PL, et al. Mammalian BTBD12 (SLX4) protects against genomic instability during mammalian spermatogenesis. PLoS Genet. 2011;7(6):e1002094.

18. Reinholdt L, Ashley T, Schimenti J, Shima N. Forward genetic screens for meiotic and mitotic recombination-defective mutants in mice. Methods Mol Biol. 2004;262:87–107.

19. Dertinger SD, Torous DK, Tometsko KR. Simple and reliable enumeration of micronucleated reticulocytes with a single-laser flow cytometer. Mutat Res. 1996;371(3-4):283–92.

20. Balmus G, Karp NA, Ng BL, Jackson SP, Adams DJ, McIntyre RE. A high-throughput in vivo micronucleus assay for genome instability screening in mice. Nat Protoc. 2015;10(1):205–15.

21. Pruitt SC, Qin M, Wang J, Kunnev D, Freeland A. A Signature of Genomic Instability Resulting from Deficient Replication Licensing. PLoS Genet. 2017;13(1):e1006547.

22. Shima N, Alcaraz A, Liachko I, Buske TR, Andrews CA, Munroe RJ, et al. A viable allele of Mcm4 causes chromosome instability and mammary adenocarcinomas in mice. Nat Genet. 2007;39(1):93–8.

23. Koren A, Massey DJ, Bracci AN. TIGER: inferring DNA replication timing from wholegenome sequence data. Bioinformatics. 2021.

24. McIntyre RE, Nicod J, Robles-Espinoza CD, Maciejowski J, Cai N, Hill J, et al. A Genome-Wide Association Study for Regulators of Micronucleus Formation in Mice. G3 (Bethesda). 2016;6(8):2343–54.

25. Maccaroni K, Balzano E, Mirimao F, Giunta S, Pelliccia F. Impaired Replication Timing Promotes Tissue-Specific Expression of Common Fragile Sites. Genes (Basel). 2020;11(3).

26. Waters CE, Saldivar JC, Hosseini SA, Huebner K. The FHIT gene product: tumor suppressor and genome “caretaker”. Cell Mol Life Sci. 2014;71(23):4577–87.

27. McAvoy S, Ganapathiraju SC, Ducharme-Smith AL, Pritchett JR, Kosari F, Perez DS, et al. Non-random inactivation of large common fragile site genes in different cancers. Cytogenet Genome Res. 2007;118(2-4):260–9.

28. Yehuda Y, Blumenfeld B, Mayorek N, Makedonski K, Vardi O, Cohen-Daniel L, et al. Germline DNA replication timing shapes mammalian genome composition. Nucleic Acids Res. 2018;46(16):8299–310.

29. Palmerola KL, Amrane S, De Los Angeles A, Xu S, Zuccaro MV, Massey DJ, et al. DNA breaks due to replication stress limit the developmental potential of human preimplantation embryos. https://papersssrncom/sol3/paperscfm?abstract_id=3825160. 2021.

30. Laks E, McPherson A, Zahn H, Lai D, Steif A, Brimhall J, et al. Clonal Decomposition and DNA Replication States Defined by Scaled Single-Cell Genome Sequencing. Cell. 2019;179(5):1207–21 e22.

31. Yin Y, Jiang Y, Lam KG, Berletch JB, Disteche CM, Noble WS, et al. High-Throughput Single-Cell Sequencing with Linear Amplification. Mol Cell. 2019;76(4):676–90 e10.

32. Minussi DC, Nicholson MD, Ye H, Davis A, Wang K, Baker T, et al. Breast tumours maintain a reservoir of subclonal diversity during expansion. Nature. 2021.

33. Gonzalez V, Natarajan S, Xia Y, Klein D, Carter R, Pang Y, et al. Accurate Genomic Variant Detection in Single Cells with Primary Template-Directed Amplification. BioRxiv. 2020.

34. Massey DJ, Koren A. High-throughput analysis of DNA replication in single human cells reveals constrained variability in the location and timing of replication initiation. BioRxiv. 2021;www.biorxiv.org/content/10.1101/2021.05.14.443897v1?ct=

35. Dertinger SD, Torous DK, Hall NE, Murante FG, Gleason SE, Miller RK, et al. Enumeration of micronucleated CD71-positive human reticulocytes with a single-laser flow cytometer. Mutat Res. 2002;515(1-2):3–14.

