## Supplementary Information for "Sequencing Micronuclei Reveals the Landscape of Chromosomal Instability"

| **Strain** | **%NCE-MN** |
| --- | --- |
| FVB/B6 male | 0.2 |
| *Rad9a^SA^* male repetition 1 | 0.9 |
| *Rad9a^SA^* male repetition 2 | 2.8 |
| *Rad9a^SA^* female | 1.56 |
| C3H female | 0.21 |
| *Chaos3* female | 4.37 |
| *Chaos3* male | 4.4 |

Table S1. Rate of micronucleation in each mouse strain.

Table S2. All raw, slope and segment data (xls file).

Sheet 1 provides the raw DNA copy number data, after filtering, for all samples. Sheet 2 provides the corresponding slope value calculated for each window. Sheet 3 provides the discrete segments called in each sample, each having a unique slope and slope value jump from the previous segment.

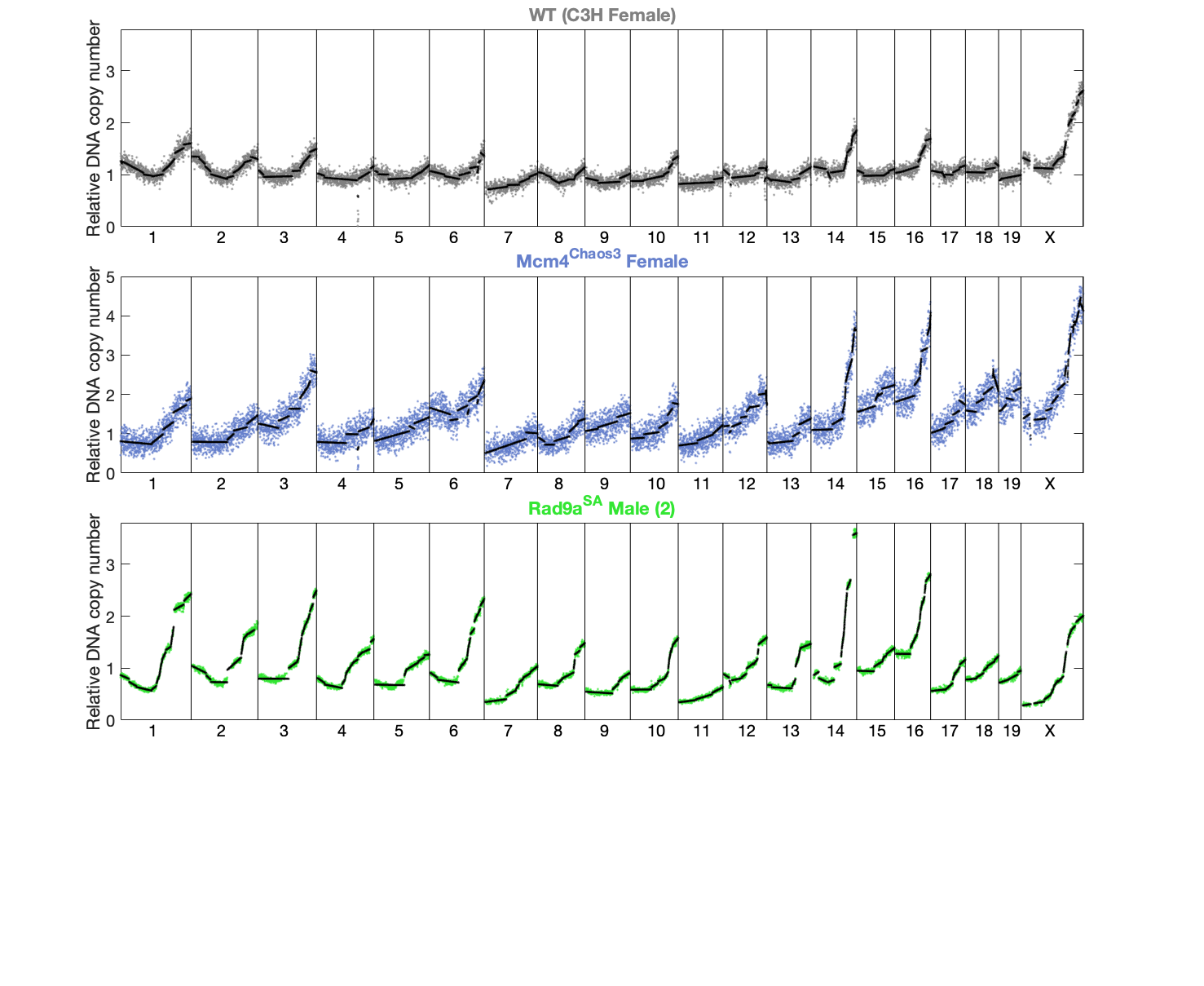

**Figure S1. MN DNA across the remaining strains (those not shown in Figure 2).**

As in Figure 2. R*ad9a^SA^* is a biological repetition of the sample shown in Figure 2. C3H is the genetic background of the *Mcm4^Chaos3^* strain.

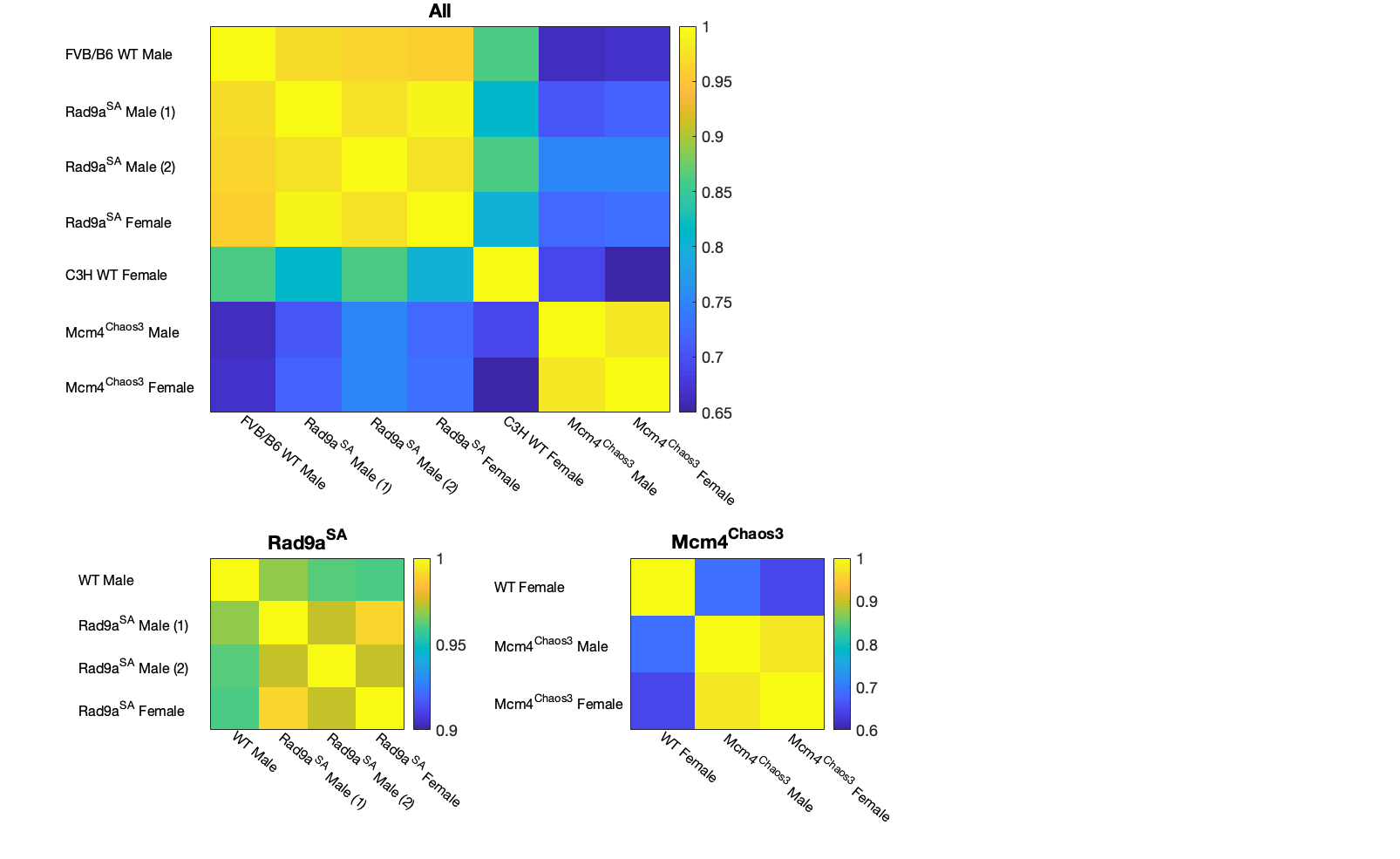

**Figure S2. Correlations of MN fragment copy number slope data for all samples.**

Top image shows all samples, bottom images are stratified by GIN models. While the experiment is highly reproducible, both genetic background and mutant status influence the landscape of chromosomal fragility as observed by MN-seq.

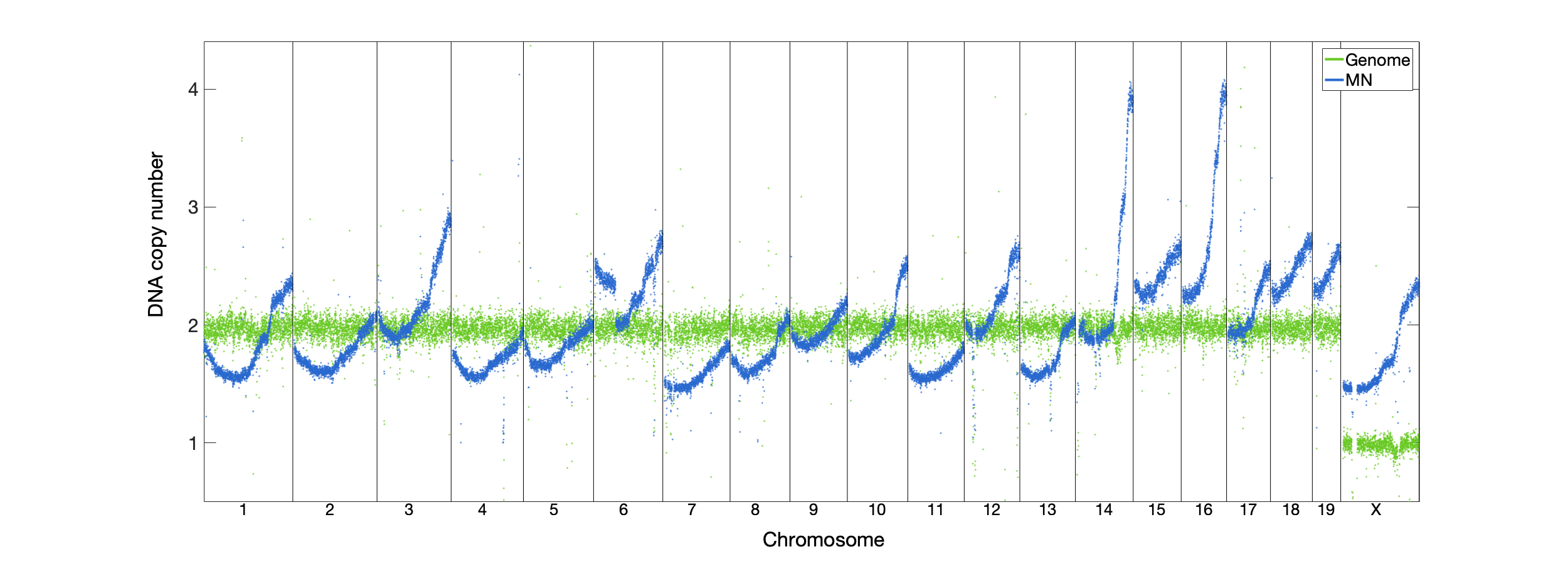

**Figure S3. MN DNA copy number is not biased by copy number variation or aneuploidy in the genome.**

MN-seq data for *Mcm4^Chaos3^* male (arbitrary units) is shown alongside whole genome sequence data for male MEF cells of the same strain.

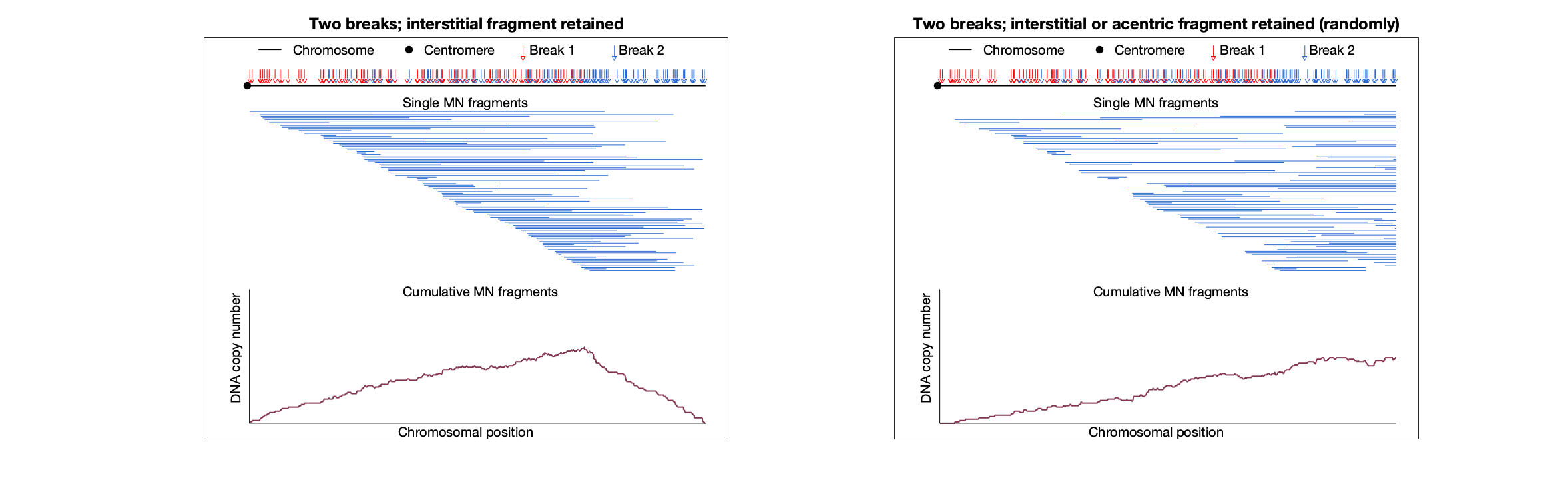

**Figure S4. Simulations of micronuclei resulting from two breaks occurring on the same chromosome.**

As in Figure 3, with two breaks on the same chromosome. The first break was simulated to occur anywhere in the leftmost ¾ of the chromosome and is indicated with a red arrow. The second break was simulated anywhere between the first break and the right end of the chromosome and is indicated with a blue arrow. The figures do not show which blue arrow is paired with which red arrow, but the single MN fragments represent the result of the two breaks. The left plot simulates a scenario in which the interstitial fragment (between the two breaks) is always the fragment retained within MN, while in the right plot, the retained fragment is randomly picked to either be the interstitial fragment or the fragment from the second break to the right end of the chromosome.

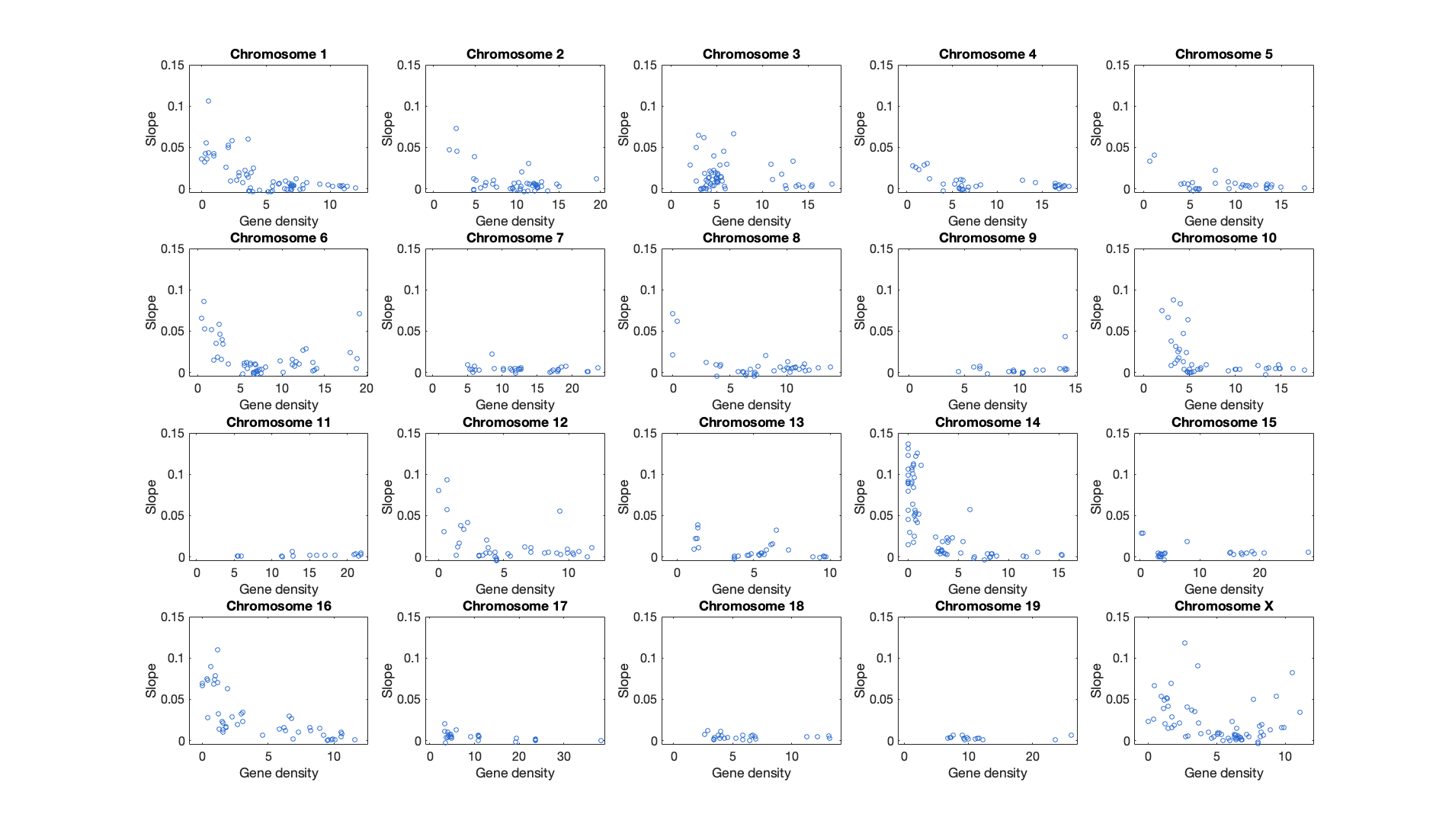

**Figure S5. Segment slope as a function of gene density, per chromosome.**

As in Figure 6A. Note that the number of segments is different between chromosomes.
